# Unitary Structure of Palindromes in DNA

**DOI:** 10.1101/2021.07.21.453288

**Authors:** Mehmet Ali Tibatan, Mustafa Sarisaman

**Affiliations:** Department of Biotechnology, Istanbul University, 34134, Vezneciler, Istanbul, Turkey; Department of Physics, Istanbul University, 34134, Vezneciler, Istanbul, Turkey

**Author notes:** Electronic address (Mustafa Sarisaman). PACS numbers: 03.65.−w, 02.30.Tb, 02.90.+p, 87.10.+e,87.14.Gg, 87.14.G−.

## Abstract

We investigate the quantum behavior encountered in palindromes within DNA structure. In particular, we reveal the unitary structure of usual palindromic sequences found in genomic DNAs of all living organisms, using the Schwinger’s approach. We clearly demonstrate the role played by palindromic configurations with special emphasis on physical symmetries, in particular subsymmetries of unitary structure. We unveil the prominence of unitary structure in palindromic sequences in the sense that vitally significant information endowed within DNA could be transformed unchangeably in the process of transcription. We introduce a new symmetry relation, namely purine-purine or pyrimidine-pyrimidine symmetries (p-symmetry) in addition to the already known symmetry relation of purine-pyrimidine symmetries (pp-symmetry) given by Chargaff’s rule. Therefore, important vital functions of a living organisms are protected by means of these symmetric features. It is understood that higher order palindromic sequences could be generated in terms of the basis of the highest prime numbers that make up the palindrome sequence number. We propose that violation of this unitary structure of palindromic sequences by means of our proposed symmetries leads to a mutation in DNA, which could offer a new perspective in the scientific studies on the origin and cause of mutation.

## INTRODUCTION

Sequence specific functionality drives the proteins enhanced functions and play role as a guidance but the question of why DNA require palindromic sequences has not clear answer yet but there are only few examples in the literature on how palindromic sequences works. Nevertheless, question of ‘are there any reason to determine the palindromic sequences as important and significant rather than arbitrary nucleic acid sequences?” might require more detailed answers.

Ordinarily, nucleic bases can have different properties from each other despite their structural similarities (See, Fig.1). For instance, adenine shows different aminoacid residue preferences than the guanine [1] or cytosine can present unique UV-absorption spectrum which differs from absorption spectrum of the other bases [2], additionally, guanine has the most reactive site in the DNA against reactive oxygen species [3]. Different aminoacid residue preference of the bases can be a defining feature by means of protein interaction with relative gene regions. Although nucleic basis have similar affinity to protons which directly affect the tautomeric stabilization levels, their substrate binding energetics may differ [4]. In other words, structural similarities and differences of nucleic bases provide a specific molecular recognition [1]. In this case, the molecular recognition is important in terms of information conservation because notably quadruplet and sextet palindromic sequences provide unique binding motifs especially for the transcription starter proteins or enzymes which can break the DNA strands.

**FIG. 1:**
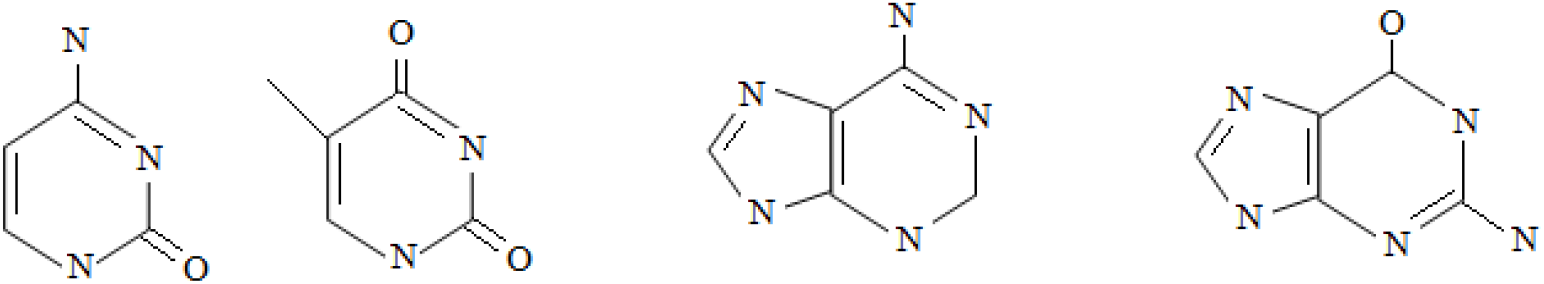
Simple ring structure of nucleobases which forms the DNA, hydrogen bonds not shown. Adenine and guanine (purines) have two-ring structure differently from cytosine and thymine (pyrimidine). Explicit unitary symmetries are displayed between adenine and thymine, and cytosine and guanine, which reflect the Chargaff’s rule.

Palindromic structure of DNA is nucleic acid sequences which are symmetrical with their sense and anti-sense strand. These specific sequences are suggested to enhance affinity for transcriptional regions [5] and more importantly they can involve the recombination repair mechanism because both of sense and anti-sense regions of palindromic DNA sequences are symmetrical which supports the enzymatic proteins to bind both strands at the same time. Studies show that palindromic sequences might help to homologous recombination and palindrome containing DNA regions are indented to replicate easier [6]. Moreover, previous studies have shown that these palindromic oligonucleotides have remarkable immunogenetic properties and potantiate the induction of interferon release from natural killer cells [7, 8]. However, above all the explanations regarding the functionality of palindrome, the best approach may be the ability of these specific sequences to form hairpin [9]. Nag and Kurst revealed that palindromic sequences present breakpoints during the cellular process which requires double-strand breaks such as meiosis [10]. Briefly, it has been suggested that palindromic sequences help DNA’s folding process (forming hairpin structures) and recognition by nucleases, as well as that RNA viruses use palindromic sequences to dimerize the viral genome, which provides protection against enzymatic degradation by the host cell. Even, it has been noted that they cause some problems with sequencing technologies due to their tendency to fold [10–12].

However, despite all these alternative explanations, the role of palindromic sequences within our genome requires more detailed explanation because it is not clear yet how energetic functionality of such symmetric structures takes advantage through the cellular mechanism, and even how they potentially differ from the non-palindromic sequences. Therefore, the roles of palindromes should be more comprehensively uncovered in order to advance nucleobase function within the DNA. Nucleotide functions and sequence-specific protein interaction has been extensively studied, and such studies are still ongoing [13]. The electrochemical properties of the nucleobases shown in Fig. 1 may explain their functionality [12–15], but this may not be sufficient to elucidate their selective function in DNA during cellular activities.

How enzymes recognize the double strand of the palindrome, why nucleobases tend to form hairpin structure during replication, and how nucleobase alignment affects DNA stability may have an electrochemical explanation. However, the study of nucleobase behavior and thus the palindromic sequence mechanism at the quantum level will improve all these explanations and provide some insight into the energetic principles of the nucleic acid sequences [13, 14]. Therefore, we present a rudimental approach to DNA at the quantum level by revealing the palindromic sequences to better understand the nature of nucleobases, which may provide some explanation for their energetic mechanism.

In view of all these fact, we present an alternative perspective to palindromes in DNA, which reflect their unitary structure distinctively. This approach leads to the emergence of unknown symmetries of palindromes with very important functions. We propose a novel symmetry relation which has not been realized yet, which gives rise to other vitally important accompanying palindromes. In particular, we focused on specific palindromes whose properties and roles are known extensively, such as **CACGTG** depicted in Fig. 2 which is called as E-box consensus, and showed the importance of the symmetry relations we found. This palindromic sequence is normally found at the protein binding site in the breast cancer cells [15–17]. For example, since human c-myc protein has been reported to be over-expressed in tumor progression, it is responsible for cancer development [18, 19]. Additionally, E-box elements within the genome provide selectivity for transcription factors to find their target sites, so these sequences are important to bring DNA recognition specificity [16]. We propose a quantum-mechanical explanation of nucleobase symmetry in order to identify the role of a palindromic site to bind these tumorigenic proteins.

**FIG. 2:**
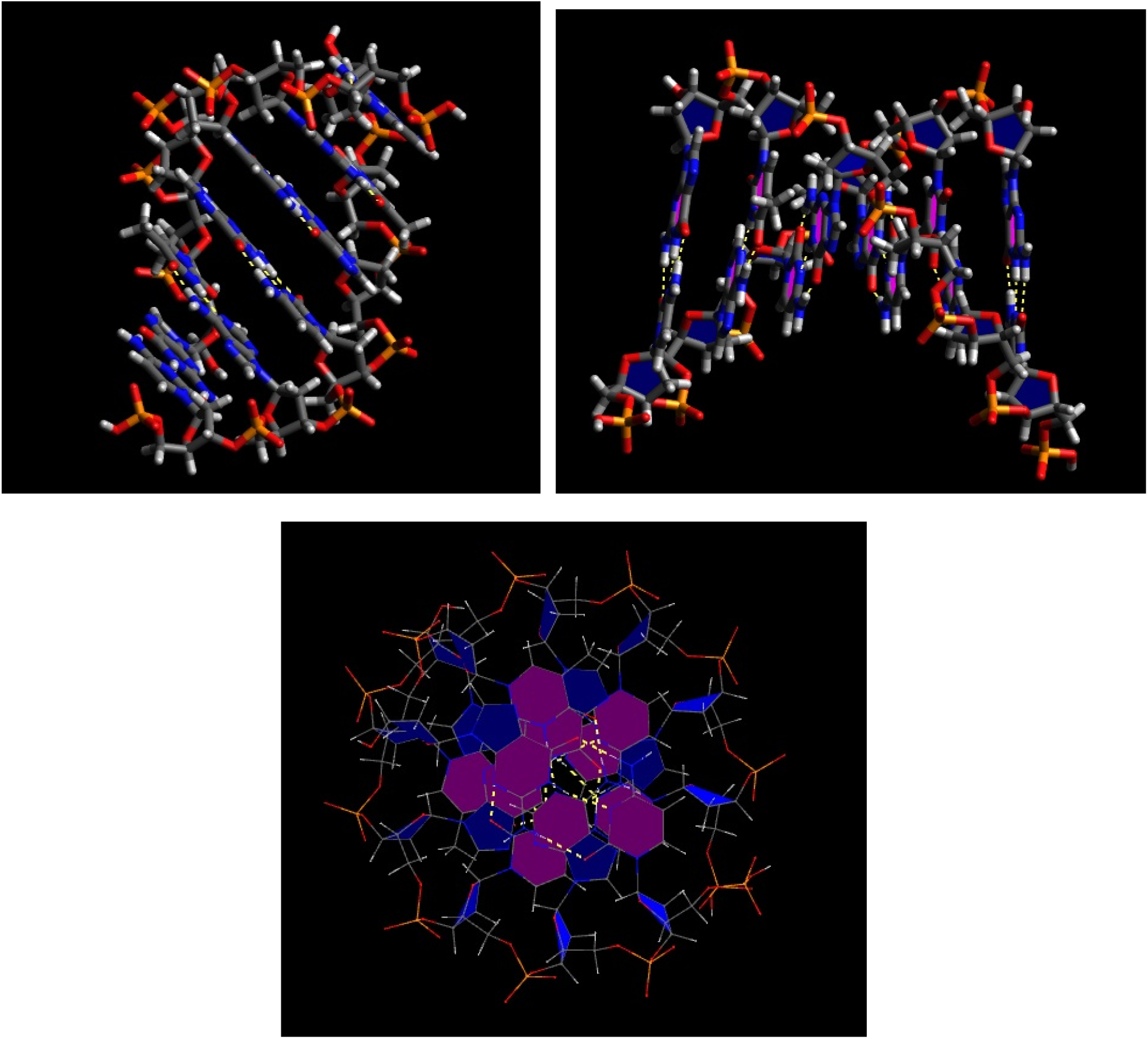
(Color online) Ball and stick view of the palindromic E-box consensus (**CACGTG** sequence). Dashed yellow lines indicate H-bonds between the nucleobases. This figure clearly shows the intrinsic unitary structure of **CACGTG** in the form of chiral symmetry of Dirac. Image prepared by using Avogadro software (ver. 1.2.0)

## PALINDROMIC STRUCTURE OF DNA AND ITS PHYSICAL SYMMETRIES

Palindromic nature of DNA is rather significant for the maintenance of vital functions of living mechanisms. This is why we persistently witness some distinguished palindromic forms in almost all important genes. To understand its vital role when examined more closely from a physical perspective, we notice that some featured physical symmetries are encountered in these particular regions of DNA. In the process of DNA replication by a specific enzyme, initially given chain of DNA is transformed into its counterpart by means of nucleotide rules (known as Chargaff’s rule). This procedure permits an inversion symmetry in the whole DNA entity. This in in fact very similar to chiral symmetries of fermions in the chiral representation of Dirac type particles in physics, in which a right-handed particle exhibits similar behavior as the left-handed one [20–22]. Fig. 3 displays the inversion symmetries in **TATA**-box sequence and **CACGTG** sequence in E-box, which reflects the chiral nature of screened palindromes.

**FIG. 3:**
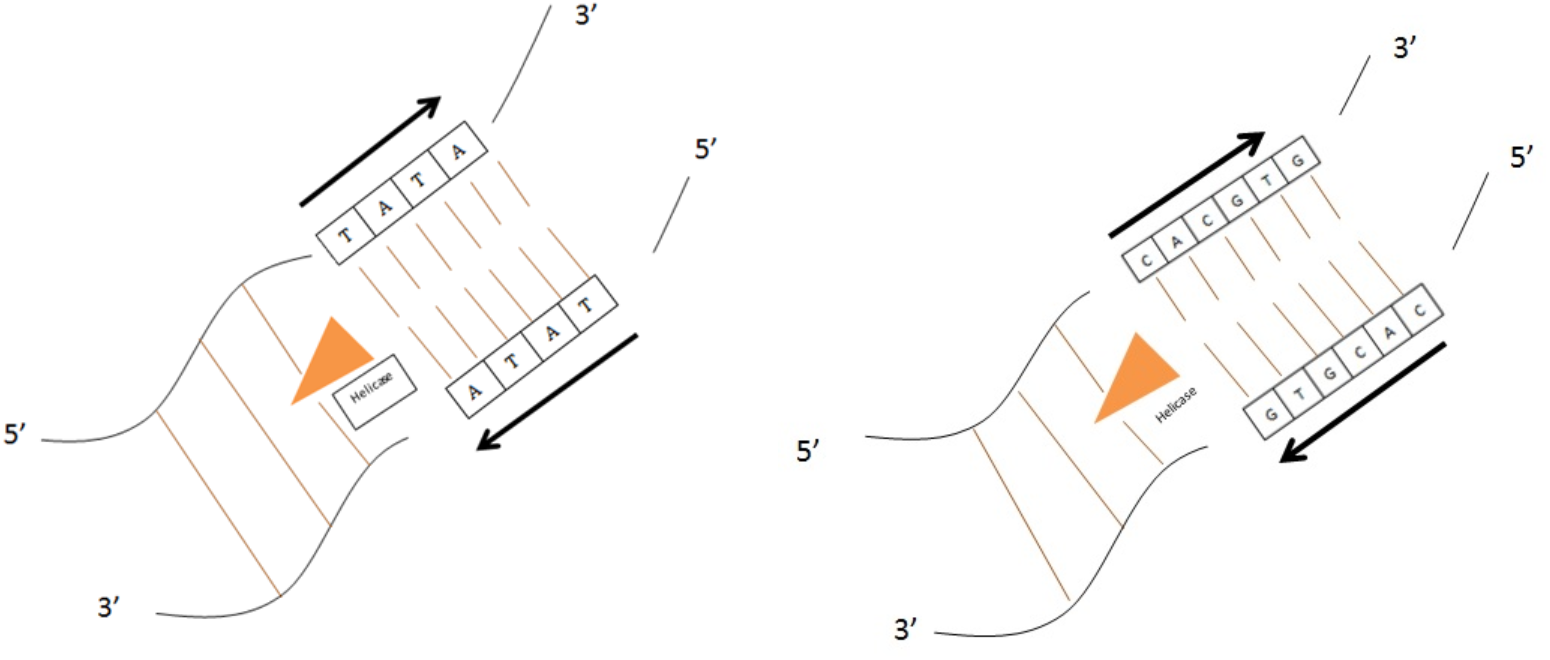
(Color online) Schematic presentation of palindromic form of **TATA**-box (left panel) and E-box sequence **CACGTG** (right panel), respectively. In both cases, Helicase enzyme is breaking hydrogen bonds between nucleotides during DNA replication or transcription to associate the polymerase enzyme activity.

**TATA**-box element within DNA is another example of palindromic sequence. It is very common and well conserved genomic sequence and can be found in many species from plants to human genome. Normally, **TATA** sequence has many different variance, but **TATA** content (thymine - adenine -adenine thymine-adenine core) is constant and giving the **TATA**-box name. Primary function of the **TATA**-box is to provide protein binding region for the transcriptional process and it locates near the promoter region of the related genes. **TATA** sequence itself is short palindromic sequence and also it is core for the consensus sequence and has many variance [23]. As for all the other palindromic sequences, **TATA** sequence has also core properties and conserved sequence within the eukaryotic cells. Thus it is another significant example to reveal palindromic sequence importance in eukaryotic genome. Moreover it can contribute the viral contamination by helping the viral transcriptional protein involvement to the genome [24, 32].

## PALINDROMES IN DNA AND THEIR UNITARY STRUCTURE

Consider a set of quantum states which consist of singlet structures |*ζ*_*i*_⟩, where the index *i* denotes the order of singlets in the DNA sequence. In principle, *i* = (1, 2). Now, consider another set of quantum states |*ξ*_*j*_⟩, which is different from the first one in basic aspects. We describe *|ζ*_1_⟩ and *|ζ*_2_⟩ by the nucleotides adenine |**A**⟩ and thymine |**T**⟩ respectively in DNA, whereas |*ξ*_1_⟩ and |*ξ*_2_⟩ represent the nucleotides guanine |**G**⟩ and cytosine |**C**⟩ respectively, see Fig. 1 for the structures of these nucleotides, see [25–31] for further details on quantum mechanical description of states.

We define an operator 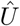 on the set |*ζ_i_*⟩ by the map 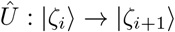 mod2. Likewise, an operator 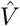 is defined on the set |*ξ_i_*⟩ by the map 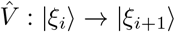 mod2. Notice that our sets of quantum states generate a periodic structure such that they form the following closed operations,

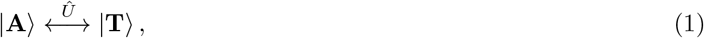

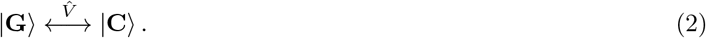

In fact, the operators 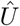 and 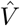 translating quantum states within a nucleotide set correspond to an enzyme, and in principle the same enzyme operates for each nucleotide pair, it operates uncustomarily on distinct nucleotide pairs. Physically, this amounts to the noncommutativity of the operators 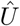 and 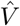, i.e. 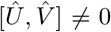. Notice that 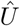 and 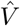 operators satisfy the following identity in a doubly acted accompanying singlet nucleotide system,

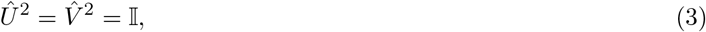

where 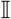 denotes the identity matrix such that our set is complete. This is in fact our completeness relation. This relation tells us that eigenvalues for the 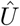 and 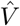 operators are given by 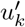 and 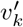

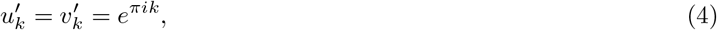

where *k* = 1, 2 gives rise to the relevant eigenvalues. We realize that eigenvalues are 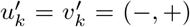. Notice that both accompanying states |**A**⟩ − |**T**⟩ and |**G**⟩ − |**C**⟩ play the role of |+⟩ and |−⟩ states in case of spin states of Dirac particles. But, we understand that the action is performed by the Pauli spin matrices 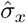 and 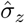 in two dimensions such that the role of the corresponding enzyme is to switch the polarity (spin) of the nucleotides. This can be seen explicitly in Fig. (1). Hence, it is noted that |**A**⟩ and |**T**⟩ states are the eigenstates of 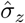, whereas |**G**⟩ and |**C**⟩ states are the eigenstates of 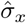 operator. Furthermore, 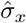 operator transforms |**A**⟩ and |**T**⟩ states into each other, just as 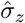 operator transforms |**G**⟩ and |**C**⟩ states into one another in a similar manner. These mutually periodic actions can be expressed in a clear way as follows,

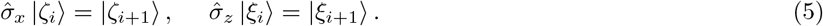

We can briefly demonstrate all these actions in the following compact forms

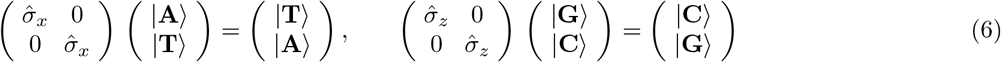

In general, we would like to find the eigenstates of the operators 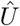 and 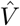 with eigenvalues 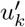 and 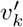, respectively. We realize that they are figured out to be the antisymmetric ordering of the states |*u_i_*⟩ and |*v_i_*⟩ such that |*u*_+_⟩ := |**G**⟩, |*u*_−_⟩ := |**C**⟩, |*v*_+_⟩ := |**A**⟩ and |*v*_−_⟩ := |**T**⟩. Thus, we observe that

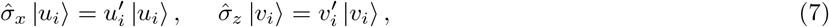

with corresponding eigenvalues 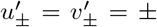. It is obvious that only way to achieve these relations is to identify 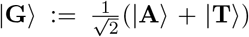 and 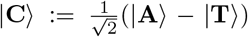, together with their counterparts 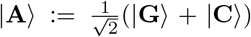 and 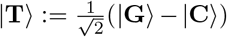. It is not difficult to observe the commutation relation of the operators 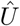 and 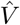, which is given by

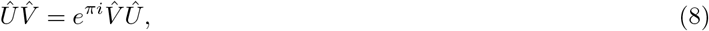

which gives rise to usual commutation relation of Pauli matrices: 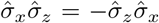. Commutation of the operators 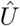 and 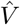 leads to the multi-ordering of nucleotides such that more complicated structures can be constructed in a given region of a DNA sequence. For example, a sequence involving two nucleotide sets can be fulfilled. Full set of such a binary set contains exactly 16 sequences[45]. We will not consider non-palindromic cases in this work, but will focus our attention to the palindromic cases which include only 4 members, which are |**AT**⟩, |**TA**⟩, |**GC**⟩, |**CG**⟩. These binary sets can be found in many DNA sequences. Notice that this set of binary palindromic structures are closed and admits a symmetry under the exchange of **A** ⇄ **T** and **G** ⇄ **C**. This closure property is inherent to the palindromic feature. Table I shows the number of palindromic sequences in binary structures.

**TABLE I:**
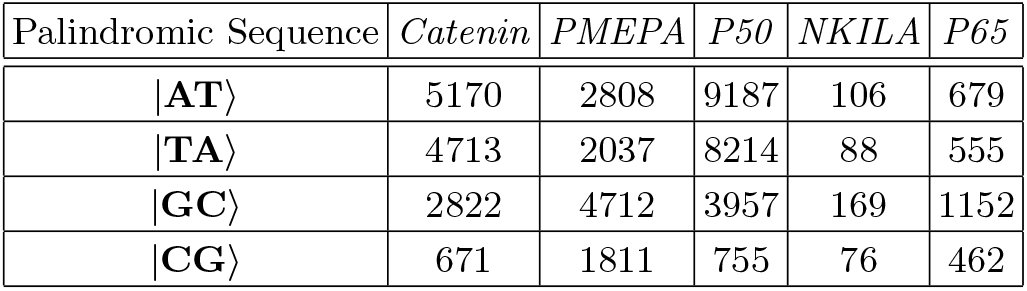
Table depicts the numbers of binary palindromic sequences in special genes of *Catenin*, *PMEPA*, *P50*, *NKILA* and *P65*. These values are extracted from FASTA sequences obtained through NCBI database.

In any palindromic sequence consisting of even multiples of binary sequence structures, one can safely choose individual states |*ζ*_*i*_⟩ and |*ξ*_*i*_⟩ as the basis of more complicated patterns. In case of a binary palindromic system, we can represent basis as |*ζ*_*i*_, *ζ*_*i*+1_⟩ := |*ζ*_*i*_⟩ ⊗ |*ζ*_*i*+1_⟩ and |*ξ*_*i*_, *ξ*_*i*+1_⟩ := |*ξ*_*i*_⟩ ⊗ |*ξ*_*i*+1_⟩. More clearly, these states are expressed as follows

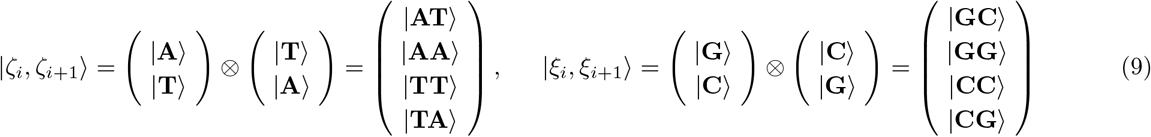

If we represent the operators transforming a purine and a pyrimidine states into their corresponding palindromic counterparts by 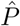 and 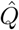 respectively, we can demonstrate that

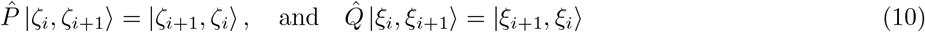

Thus, we can show that 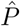 operator can be written as 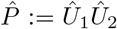 for the pure purine state |*ζ*_*i*_, *ζ*_*i*+1_⟩ and 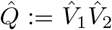 for the pure pyrimidine state |*ξ*_*i*_, *ξ*_*i*+1_⟩. Here, subindices 1 and 2 represent the order of the operator that acts on the binary state in the corresponding order. Since the same enzyme operates distinctively between two sequences, we may describe that 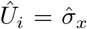 and 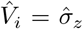, and therefore 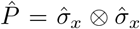 and 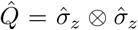. Tensor product implies the ordered operator actions. In particular, one can show Eq. 10 in matrix notation as follows

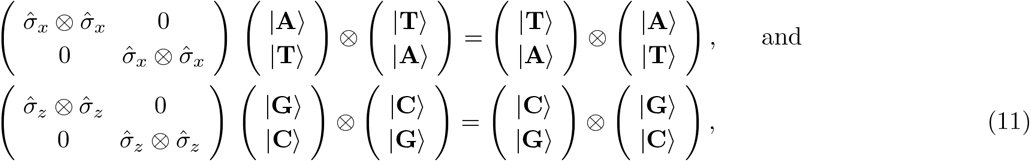

or, more explicitly,

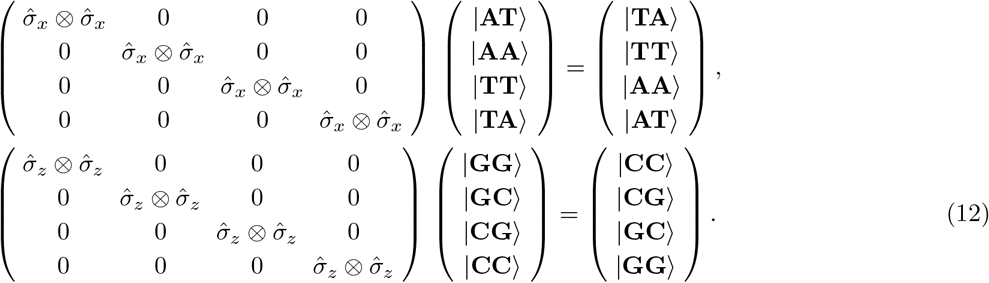

Notice that eigenvalues of 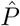 operator is given by *e*^*iπ*(2*k*+1)^ for *k* = 1, 2 mod 2. But, since *k* is an integer, this corresponds to −1 always.

Likewise, one can continue with quadruple and sextet structures. Notice that one needs to build appropriate basis for the vector space of relevant DNA sequence. In case of quadruple structure, there 256 distinct species. But not all of them are of palindromic nature. Palindromic cases are |**AATT**⟩, |**ATAT**⟩, |**AGCT**⟩, |**ACGT**⟩, |**TTAA**⟩, |**TATA**⟩, |**TGCA**⟩, |**TCGA**⟩, |**CATG**⟩, |**CTAG**⟩, |**CCGG**⟩, |**CGCG**⟩, |**GATC**⟩, |**GTAC**⟩, |**GGCC**⟩, |**GCGC**⟩, which can be figured out to be |*ζ*_*i*_, *ζ*_*i*_, *ζ*_*i*+1_, *ζ*_*i*+1_⟩, |*ζ*_*i*_, *ζ*_*i*+1_, *ζ*_*i*_, *ζ*_*i*+1_⟩, |*ζ*_*i*_, *ξ*_*i*_, *ξ*_*i*+1_, *ζ*_*i*+1_⟩, |*ζ*_*i*_, *ξ*_*i*+1_, *ξ*_*i*_, *ζ*_*i*+1_⟩, |*ξ*_*i*_, *ξ*_*i*_, *ξ*_*i*+1_, *ξ*_*i*+1_⟩, |*ξ*_*i*_, *ξ*_*i*+1_, *ξ*_*i*_, *ξ*_*i*+1_⟩, |*ξ*_*i*_, *ζ*_*i*_, *ζ*_*i*+1_, *ξ*_*i*+1_⟩, |*ξ*_*i*_, *ζ*_*i*+1_, *ζ*_*i*_, *ξ*_*i*+1_⟩. Notice that there are two types of symmetry in this palindromic set: first one is among accompanied purines and pyrimidines which are specified to be *ζ*_*i*_ ↔ *ζ*_*i*+1_ and *ξ*_*i*_ ↔ *ξ*_*i*+1_. Second one is among purines and pyrimidines in mixed forms, which is simply provided by *ζ*_*i*_ ↔ *ξ*_*i*_. These symmetries are peculiar to the unitary feature of palindromic structures. We realize that first symmetry takes place between purine or pyrimidine structures such that certain cross mappings occurs. Hence, we simply call it a *pp-symmetry* (purine-pyrimidine symmetry). pp-symmetry is the natural symmetry that occurs usually in the any enzyme activity in transcription process of DNA. On the other hand, second symmetry relation is new and requires mapping a purine or pyrimidine phases into a its corresponding counterpart within the same phase, i.e. **A** ↔ **G** and **T** ↔ **C**, that is why we call it *p-symmetry* (here, p implies purine or pyrimidine, i.e, symmetry of purine-purine or pyrimidine-pyrimidine nucleotides). p-symmetry is distinguished as a result of unitary structure of DNA such that important palindromic features can be attained because of this emergent symmetry relation. For instance, **TATA**-box has the accompanying pp-symmetric partner **ATAT** and p-symmetric partner **CGCG**-box, which is the minimum DNA binding element and an important palindromic sequence. The multiple **CGCG** cis-elements are found in promoters of genes such as those involved in ethylene signaling, abscisic acid signaling, and light signal perception. Diagram given in fig. 4 demonstrates the prescribed symmetry flows of singlet and doublet states.

**FIG. 4:**
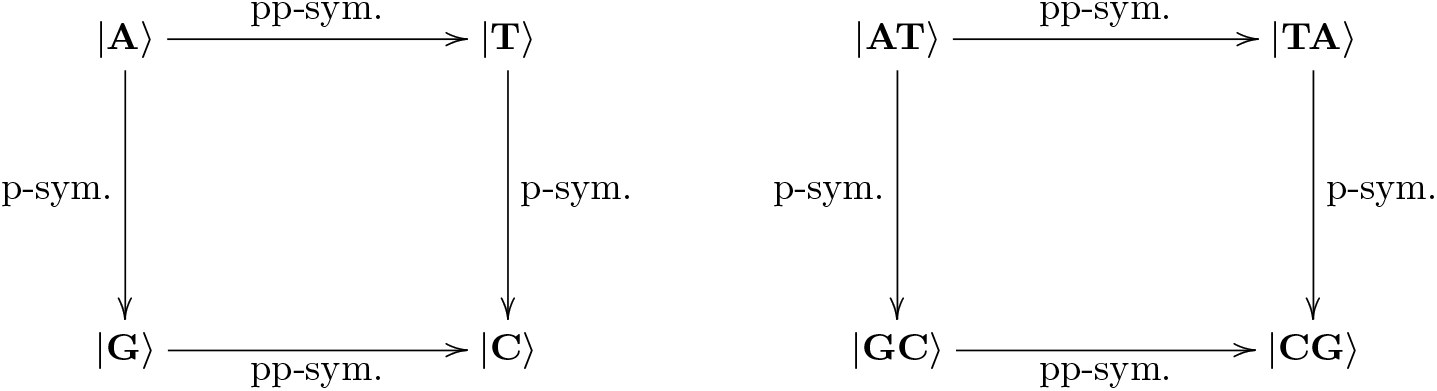
Palindromic mapping shows the flow of p- and pp-symmetries in singlet and doublet structures.

More precisely, in view of genetic perspective, it is well known that adenin transition to guanine and cytosine transition to thymine can be occurred in nature which leads to mutations. According to molecular dynamic simulations and ab inito experiments, these transitions are caused by energetic profiles of the purine and pyrimidine which are related to their different stabilities [4, 38, 39]. However, we suggest that their apparent structural similarities may be explained by means of their hidden symmetry relations as we extracted here, which are subsymmetry groups of an overall unitary symmetry in genome. In other words, repetitive palindromic sequence symmetry may be associated with adenin-guanine and thymine-cytosine structural similarities (see, Fig. 1) according to our symmetry model of unitary structure.

Moreoever, it is observed that our suggested implications can be regarded as an extension to Chargaff’s rules in which nucleic bases should have the same amount on each strand and have similar frequencies. Indeed, statistical studies approved these rules and each nucleic base has similar frequency at the opposite strand within DNA [40]. However, palindromic sequences have mirror symmetry in DNA, which is unique sequence-ordering because they present the same alignment in reverse order at the opposite strand. Thus, if we apply the Chargaff’s parity rules to our novel approach, we can implicate that palindromic sequences with a symmetry embody a more general symmetry emphasized by Chargaff’s rule. If we consider the functionality of the palindromic sequences, our proposed symmetries have purpose to controlling the conservation and/or accessibility of the genomic information, specifically help the transcriptional proteins reach the information to process [41]. With this study, we present an approach to explain the symmetrical nature of these repetitive sequences which are the part of another but more substantial symmetry in the genome of organisms. Besides, we figure out that the number of the palindromic sequence carriers (i.e., nucleic bases) can be controlled by our symmetry predictions. This provides certain palindromic structures to exist in nature, because nature allows only symmetric configurations in palindromes. As we indicate in our examples in Tables I and II, the number of palindromic sequence is drastically reduced once its sequential length increases. At that point we assume ideal conservation and accessibility of a gene region might be associated with sequential length of the palindrome. This could be related to energy considerations as well, but content of our work is beyond that level.

**TABLE II:**
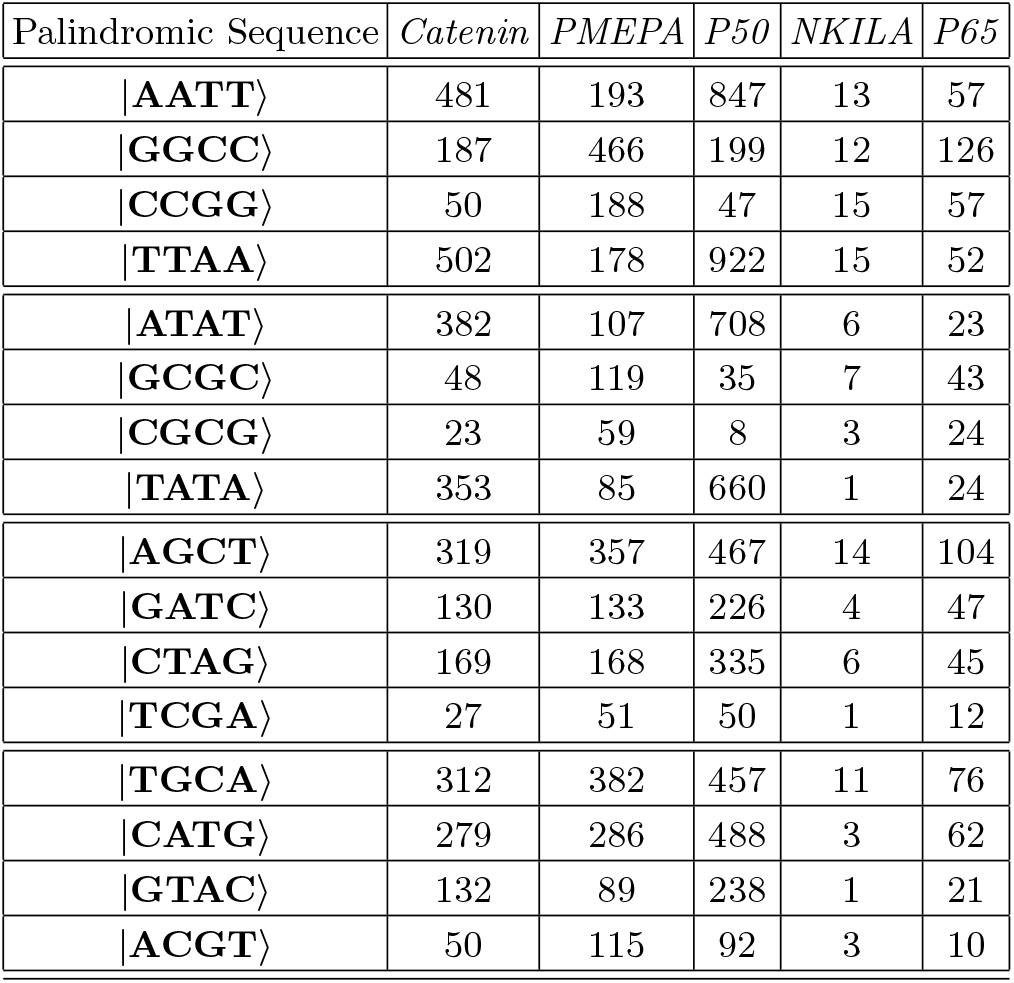
Table depicts the numbers of quadruple palindromic sequences in special genes of *Catenin*, *PMEPA*, *P50*, *NKILA* and *P65*. These values are extracted from FASTA sequences obtained through NCBI database.

Table II demonstrates the number of incidence of quadruple palindromic structures corresponding to certain types of genes. Fig. 5 shows the symmetry flows of all palindromic mappings in quartet structures. Fig. 6 demonstrates the palindromic paths for binary and quadruplo structures given in Tables I and II. Notice the similar behaviors of certain genes.

**FIG. 5:**
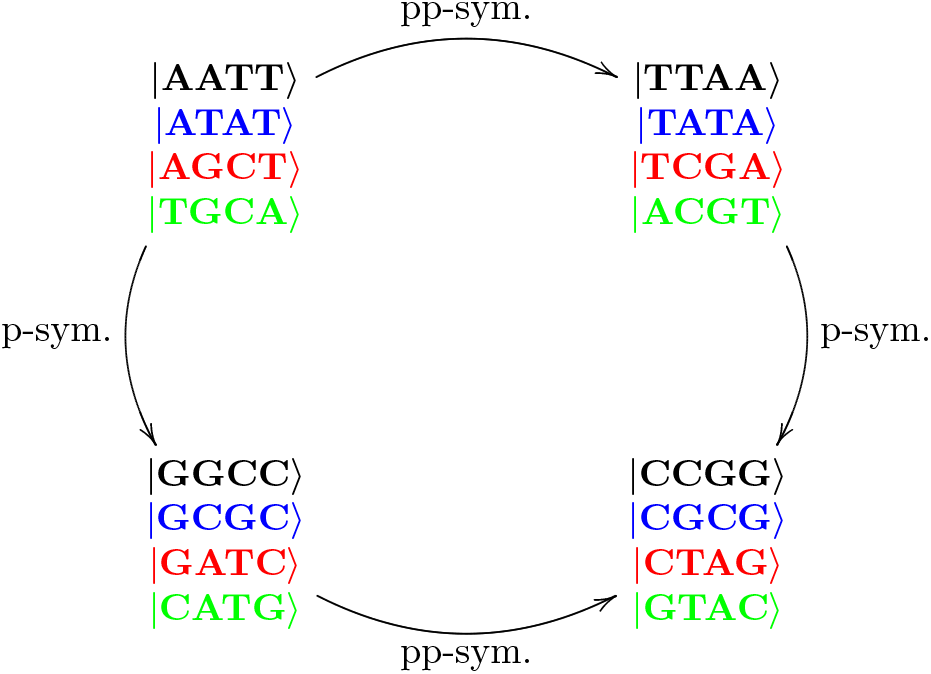
(Color Online) Diagram displays all palindromic mappings corresponding to the flow of p- and pp-symmetries in quartet structures.

**FIG. 6:**
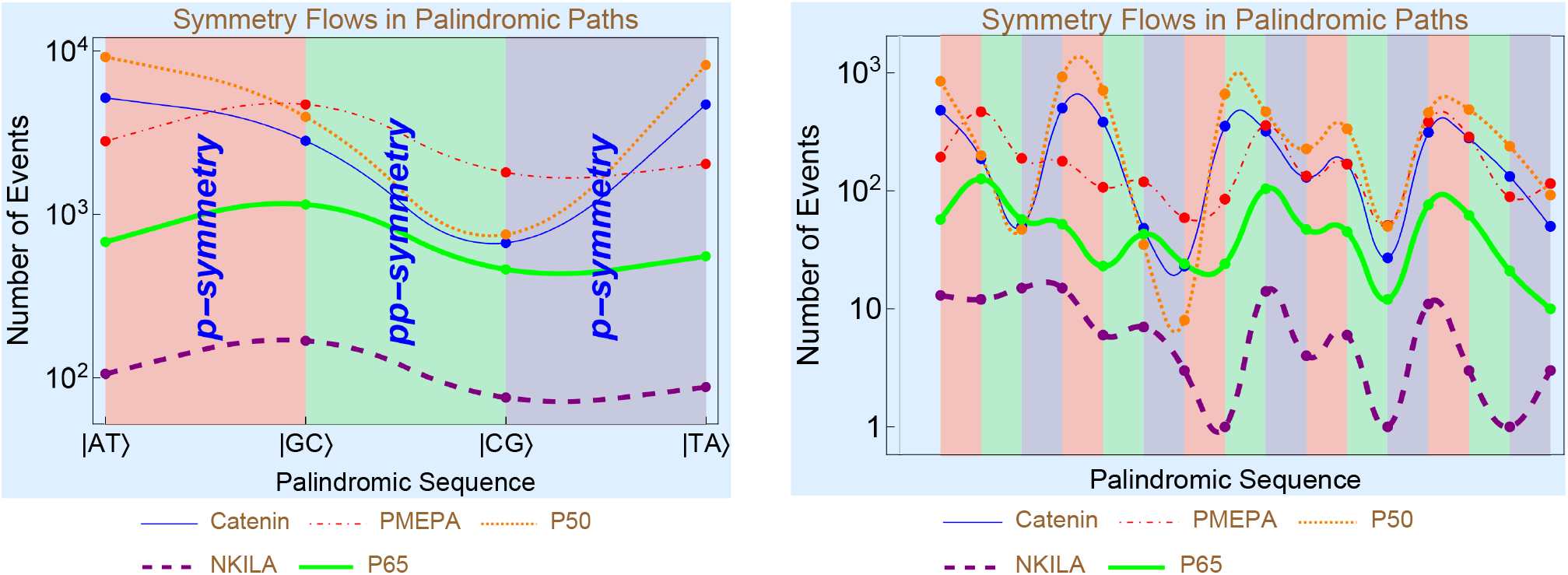
(Color online) Figures depict the palindromic paths for binary (left panel) and quadruple (right panel) structures provided in Tables I and II. Palindromic sequences and corresponding symmetry relations are not shown explicitly in quadruple structures because they fill the whole axis. In these plots, pink- and gray-colored zones correspond to p-symmetry transitions whereas green-colored zones correspond to pp-symmetry transition regions. Notice the similar behaviors of *Catenin* and *P50* genomes and that of *PMEPA*, *P50* and *NKILA*.

As can be seen from this symmetry table implying palindromic symmetries, each category contains rather crucial palindromic sequences. Another important example of short palindromic sequences in this list is 5’-**GATC**-3’ regions in the E. coli genome. Mismatch nucleotide is a type of mutation in the organisms and the mismatch repair (MMR) mechanism uses the **GATC** sequences to find and repair the damaged regions on the DNA [33]. Many enzymes taking part in this complicated molecular repair mechanism. **GATC** sequences are initiators to assemble this complex molecular machinery. First, in this repair process **GATC** sequence is methylated on the leading strand (5’ to 3’ strand) and after this **GATC** is recognized by relevant enzymatic proteins that are capable of recognizing the breakings or mismatch regions on the DNA. **GATC** sequences are the initiating points for the MMR protein repair process, through this MMR proteins can scan the DNA strand and able to find mismatch base pair then the exonucleases begin their replacement of the mutated region [34]. Additionally, **GATC** sequences repeat themselves in every 256 nucleotides through the prokaryotic genome to facilitate the MMR proteins to scan the genome more efficiently [35]. However, in humans, **GATC** may represent methylation sites to provide chromosomal stability and control the protein synthesis [36]. In other words, **GATC** sequence presents a good methylation site in the organisms and its frequency attenuates the recognition by the relevant proteins. This methylation mechanism is very important to regulate genomic information [37].

Another consequence of p- and pp-symmetries is that quadruple structures can be obtained by employing doublets via these symmetries. For instance, |*ζ*_*i*_, *ζ*_*i*_, *ζ*_*i*+1_, *ζ*_*i*+1_⟩ state can be expressed by doublets as |*ζ*_*i*_, *ζ*_*i*_, *ζ*_*i*+1_, *ζ*_*i*+1_⟩ = |*ζ*_*i*_, *ζ*_*i*_⟩ ⊗ |*ζ*_*i*+1_, *ζ*_*i*+1_⟩. pp-symmetry converts this quartet into its counterpart as |*ζ*_*i*+1_, *ζ*_*i*+1_*ζ*_*i*_, *ζ*_*i*_⟩ = |*ζ*_*i*+1_, *ζ*_*i*+1_⟩ ⊗ |*ζ_i_, *ζ*_i_*⟩. In other words, pp-symmetry acts on doublets similar to single states, i.e. 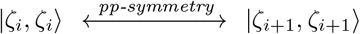 and 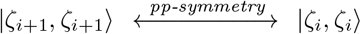. Likewise, p-symmetry acts on this state to give rise to its accompanying symmetric state: 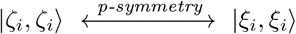 and 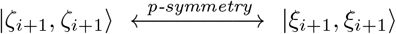. Thus, we obtain 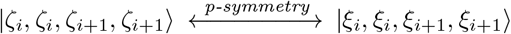 in quartet state. It is understood that one can generalize p- and pp-symmetries into doublets and higher order palindromic structures. Although there is no mixed forms of purines and pyrimidines in palindromic doublet states, quartet palindromic sequences allow mixed structures of doublet states. Therefore, in order to obtain all quartet palindromic sequences, one can consider all mixed combinations of |*ζ_i_, ζ_i_*⟩, |*ζ_i_, ξ_i_*⟩, |**ξ*_i_, ζ_i_*⟩ and |**ξ*_i_, *ξ*_i_*⟩, which yields the number of distinct categories given in Table (II). In view of these symmetry relations, it seems that operator descriptions of quartet palindromic structures are more accessible.

Sextet structures deserve a special attention because of their crucial role in the transfer of important information that ensures the continuity of cell cycle. All features in view of symmetry operations can be extended to this notable palindromic structures. One can build up a sextet system by means of individual nucleotide basis of pure |*ζ, ζ, ζ, ζ, ζ, ζ*⟩ states (i.e., |*ζ_i_, *ζ*_i_, *ζ*_i_, ζ_i+1_*, *ζ*_*i*+1_, *ζ*_*i*+1_⟩, |*ζ*_*i*_, *ζ*_*i*_, *ζ*_*i*+1_, *ζ*_*i*_, *ζ*_*i*+1_, *ζ*_*i*+1_⟩, |*ζ*_*i*_, *ζ*_*i*+1_, *ζ*_*i*+1_, *ζ*_*i*_, *ζ*_*i*_, *ζ*_*i*+1_⟩, |*ζ*_*i*+1_, *ζ*_*i*+1_, *ζ*_*i*+1_, *ζ*_*i*_, *ζ*_*i*_, *ζ*_*i*_⟩, |*ζ*_*i*+1_, *ζ*_*i*_, *ζ*_*i*_, *ζ*_*i*+1_, *ζ*_*i*+1_, *ζ*_*i*_⟩, |*ζ*_*i*_, *ζ*_*i*+1_, *ζ*_*i*_, *ζ*_*i*+1_, *ζ*_*i*_, *ζ*_*i*+1_⟩, |*ζ*_*i*+1_, *ζ*_*i*+1_, *ζ*_*i*_, *ζ*_*i*_, *ζ*_*i*_, *ζ*_*i*+1_⟩, |*ζ*_*i*+1_, *ζ*_*i*_, *ζ*_*i*+1_, *ζ*_*i*_, *ζ*_*i*+1_, *ζ*_*i*_⟩), pure **ξ*, *ξ*, *ξ*, *ξ*, ξ, ξ* states which are formed by the exchange of *ζ* ↔ *ξ* and mixture of these two states. It is obvious that there are 2^6^ = 64 palindromic family of states. But notice that these peculiar states could be established by employing doublet basis. All mixed combinations of doublet states give rise to construct any sextet state. Besides, one can also use triplets as the basis of sextet structures. All their combinations consist of the members of the set: |*ζ, ζ, ζ*⟩, |*ζ, ζ, ξ*⟩, |*ζ, ξ, ζ*⟩, |*ζ, ξ, ξ*⟩, |*ξ, ζ, ζ*⟩, |*ξ, ζ, ξ*⟩, |*ξ, ξ, ζ*⟩, |*ξ, ξ, ξ*⟩. Obviously, any sextet state can be expressed as mixed combinations of doublets or triplets, this is because 6 can be factorized as 6 = 2.3. But for any quartet state this is 4 = 2.2 = 2^2^. It is understood that we can obtain a formalism to construct higher order palindromic sequences by means of smaller sequences according to nucleotide basis of prime integer numbers that corresponding palindromic sequence contains. If we express the palindromic sequence by the number *N*, we can decompose it in terms of *n* prime integers as 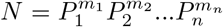, where *P_i_* is the prime integer and *m_i_* is the corresponding power of *P_i_* that is an integer. Therefore, one can describe a palindromic sequence with *N* individual basis in terms of *P_i_* sub-basis. This could be done by *n* different ways so that corresponding palindromic state preserves its unitary structure. Figure 7 shows the symmetry diagram of **CACGTG** palindrome in sextet, doublet and triplet basis.

**FIG. 7:**
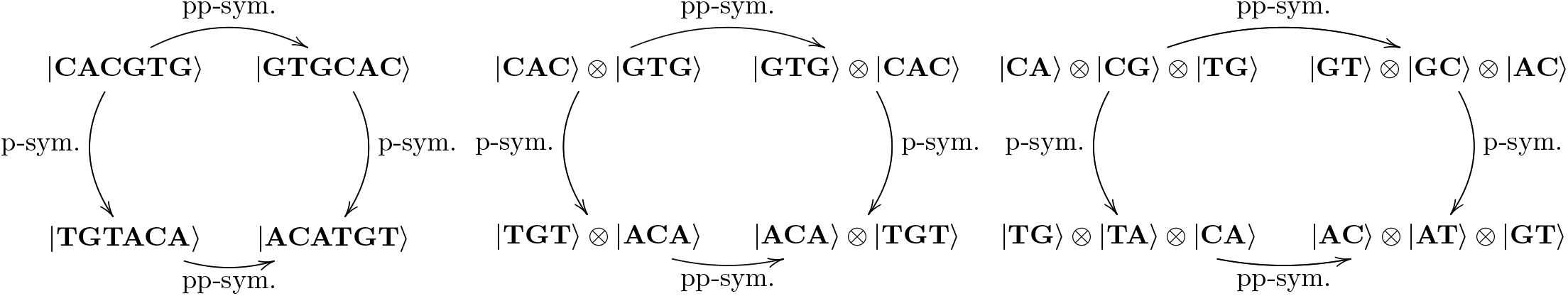
Diagram displays all palindromic mappings of **CACGTG** sequence in singlet, doublet and triplet basis corresponding to the flow of p- and pp-symmetries in sextet structures.

Notice that although one can construct higher order palindromic sequences in principle, their appearance in any genome are so rare because their probability of getting together is very low. This could be explained by energy considerations in genomic sequence. But, this requires more elaborate treatment of the subject which is beyond the scope of this work.

## CONCLUDING REMARKS

In this study, we aim to reveal the quantum effects of palindromic sequences in DNA in view of symmetry perspectives, especially unitary symmetry. Unitary symmetry basically points out the preserving important information of DNA inherent to the essence of continuity of cell cycle. Our approach exhibits the existence of subsymmetry groups of unitary symmetry in palindromes, which are called p- and pp-symmetries. pp-symmetry has been the commonly used natural symmetry of DNA known as Chargaff’s rules. However, p-symmetry is also found in palindromes, which contribute to the unitary symmetry. By means of these symmetry relations, we can better understand the structures and formations of higher order palindromes and why they get rare with the increasing number of sequences.

Dirac notations describing the basic hallmarks of the states of palindromes stipulate new qualifications of these peculiar configurations in DNA, especially in view of the quantum aspects. It is understood that subsymmetry groups of unitary structure specify the role played by palindromes. Their distinguished behaviors are strictly controlled by symmetry rules dictated by the quantum features. It is revealed that the more we understand the symmetry properties of palindromes, the more we can learn about palindromes. Our findings propose that rare number of sequences for higher order palindromes in various genes can be explained by the low probability of the bases forming palindromes to coalesce together. We find out that relevant bases of higher order palindromes can be constructed by any prime integer sequences that generate the palindrome.

Our approach in this study will help to reveal the further quantum nature of DNA. Palindromic examples we presented here are oncogenic associated gene sequences and contain many palindromic sequences in their genomic DNA regions. DNA molecules contain enormous amount of data which requires high level of conservation within and/or among the organisms to prevent mutations. Palindromic sequences may present unique features and has some advantages like easier enzymatic process. However, we suggest that palindromic sequences require more conservation or stabilization than other type of sequence and their symmetrical structures render them quintessential points in the manner of data transfer plus to enzymatic regulation and breaking points during the replicative process within cell division.

Here, we suggest symmetrical nature of these specific sequences renders the DNA more conservative in these points but any breaking or damaging this symmetry cause mostly carcinogenic or mutagenic results. As mentioned in the other sections in this study, we reveal the quadruplet and sextet palindrome sequence numbers in the *PMEPA*, *NKILA*, *P65*, *P50*, and *β*-*Catenin* genomic regions on the DNA. These molecules play important roles on the oncogenic signaling pathways and have association between them [42–44]. Quadruplet structures are more common while sextet structures more scarce. Therefore, we suggest that, longer palindromic structures have more specified roles within the coding regions on the DNA and governing the conservative significance of the sequence which it is belong.

We need to elaborate these structures by considering the mutation models which may shed light on the mutagenic and oncogenic mechanism of the mutation in point of symmetrical breaking view. Also, we need to interpret more genomic regions as much as possible in order to understand more clearly whether the palindromic sequence lengths associate with the power of conservation. Explaining the mutagenic models in Dirac notations may direct us to interpret the difference between asymmetrical and symmetrical structures of the DNA regions. Finally, with the approach we proposed in this work, we can better understand whether viral pathogenes including coronovirus has a symmetrical structure in their genome and whether they recognize palindromes in DNA easily. This can help us better understand their nature and find effective ways to avoid them.

## Acknowledgement

We acknowledge the support of Scientific Research Projects Coordination Unit of Istanbul University (IU BAP) under the project number: FDK-2017-25162.

## Author Contributions

Both authors contributed equally to the whole work including calculations, analysis and manuscript typing.

## Competing Interests

The authors declare no competing interests.

## Notes

### Competing Interest Statement

The authors have declared no competing interest.

